# Recombinant chimpanzee adenovirus AdC7 expressing dimeric tandem-repeat RBD of SARS-CoV-2 spike protein protects mice against COVID-19

**DOI:** 10.1101/2021.02.05.429860

**Authors:** Kun Xu, Yaling An, Qunlong Li, Weijin Huang, Yuxuan Han, Tianyi Zheng, Fang Fang, Hui Liu, Chuanyu Liu, Ping Gao, Senyu Xu, William J. Liu, Yuhai Bi, Youchun Wang, Dongming Zhou, Qinghan Wang, Wenli Hou, Qianfeng Xia, George F. Gao, Lianpan Dai

## Abstract

A safe and effective vaccine is urgently needed to control the unprecedented COVID-19 pandemic. Four adenovirus vectored vaccines expressing spike (S) protein have advanced into phase 3 trials, with three approved for use. Here, we generated several recombinant chimpanzee adenovirus (AdC7) vaccines expressing S, receptor-binding domain (RBD) or dimeric tandem-repeat RBD (RBD-tr2). We found vaccination via either intramuscular or intranasal route was highly immunogenic in mice to elicit both humoral and cellular (Th1-based) immune responses. AdC7-RBD-tr2 showed higher antibody responses compared with both AdC7-S and AdC7-RBD. Intranasal administration of AdC7-RBD-tr2 additionally induced mucosal immunity with neutralizing activity in bronchoalveolar lavage fluid. Either single-dose or two-dose mucosal administration of AdC7-RBD-tr2 protected mice against SARS-CoV-2 challenge, with undetectable subgenomic RNA in lung and relieved lung injury. These results support AdC7-RBD-tr2 as a promising COVID-19 vaccine candidate.

## Introduction

As of 1 February, 2021, the pandemic of COVID-19 has accounted for more than one hundred million laboratory-confirmed cases and two million deaths (*1-3*). The outbreak is still growing rapidly worldwide (*2*). Development of a safe and effective COVID-19 vaccine is urgently needed (*4*). Multiple platforms have been used to develop vaccines against COVID-19, including inactivated virus (*5-9*), live attenuated virus (*10*), protein subunit (*11-14*), virus-like particles (*15-17*), virus vectored vaccine (*18-25*), mRNA (*26-29*) and DNA(*30, 31*). Both human and chimpanzee adenovirus vectors were widely used for vaccine development. So far, four adenovirus vector vaccines against COVID-19 entered phase III clinical trials, which are ChAdOx1 nCoV-19 (University of Oxford and AstraZeneca) (*21, 32, 33*), Ad5-nCoV (Beijing Institute of Biotechnology and CanSino Biological Inc.) (*18, 19*), rAd26-S+rAd5-S (Gamaleya Research Institute) (*20*) and Ad26.COV2.S (J&J, with Janssen Pharmaceutical Companies) (*24, 34, 35*). Three of them have been approved for use. These recombinant adenovirus vaccines express full-length S protein of SARS-CoV-2. Besides, intranasal administration of a simian Ad36 vectored vaccine expressing prefusion-stabilized full-length S protein conferred mice protection and almost entirely prevented SARS-CoV-2 infections in both the upper and lower respiratory tracts (*22*).

S protein is comprised of S1 and S2 subunit, in which, the receptor binding domain (RBD) of S1 is responsible for recognizing and engaging its host cellular receptor protein angiotensin-converting enzyme 2 (ACE2), and S2 accounts for membrane fusion of virus and host cell (*36-39*). Therefore, S protein is a major target for the COVID-19 vaccine. Potent neutralizing antibodies are mainly focused on S protein RBD to block the receptor-binding (*40-52*). Previous studies about feline infectious peritonitis virus (FIPV; a feline coronavirus) found that immunization of cats with recombinant vaccinia virus expressing FIPV S protein resulted in early death after challenge with FIPV. It was thought to be caused by antibody-dependent enhancement (ADE) of FIPV infection (*53-55*). In addition, sera from animals vaccinated with SARS-CoV S protein could exacerbate virus infection *in vitro* through ADE, which posed serious safety issues for vaccine development (*56-58*). Since weakly- or non-neutralizing antibodies are believed as the cause of ADE during both flavivirus and coronavirus infections, concerns should be taken into the vaccine design (*59-61*). Since most of potent neutralizing monoclonal antibodies are against RBD of SARS-CoV-2 (*62, 63*), RBD is an attractive vaccine target. Recombinant SARS-CoV-2 RBD protein vaccines were reported to elicited high neutralizing antibodies in mice without ADE (*64, 65*). Therefore, we sought to develop COVID-19 vaccines based on both full-length S and RBD.

Here, we developed virus vectored vaccines based on chimpanzee adenovirus type 7 (AdC7), a rare serotype in the human population with the advantage of low level of pre-existing immunity (*66, 67*). We used a design of tandem repeat RBD-dimer (RBD-tr2) as antigen to increase the immunogenicity. This design has been used in our protein subunit COVID-19 vaccine that has advanced in phase 3 clinical trials (*68, 69*). Our adenovirus-based vaccines are aiming to further enhance the T cell responses. AdC7 expressing full-length or monomeric RBD were generated for comparison. Vaccination via intramuscular and intranasal routes were evaluated to dissect the systemic and mucosal immune responses, with the latter are believed to be beneficial for protection in respiratory system (*70, 71*). These results will provide crucial guidance for the further clinical trials.

## Results

We constructed three replication-incompetent AdC7 vaccines expressing SARS-CoV-2 full-length S, RBD and RBD-tr2, respectively (Fig. 1A). The expression cassettes were inserted into the E1 region of the AdC7 vector with E1 and E3 deletion. Western blot was conducted to confirm antigen expression in HEK 293T cells infected with the recombinant adenovirus. All these antigens were detected in the cell lysates (fig. S1A). Both monomeric RBD and RBD-tr2 were secreted in the culture supernatants (fig. S1B). Particularly, the antigen expression of AdC7-RBD-tr2 was much higher than AdC7-S and AdC7-RBD (fig. S1, A and B).

**Fig. 1.**
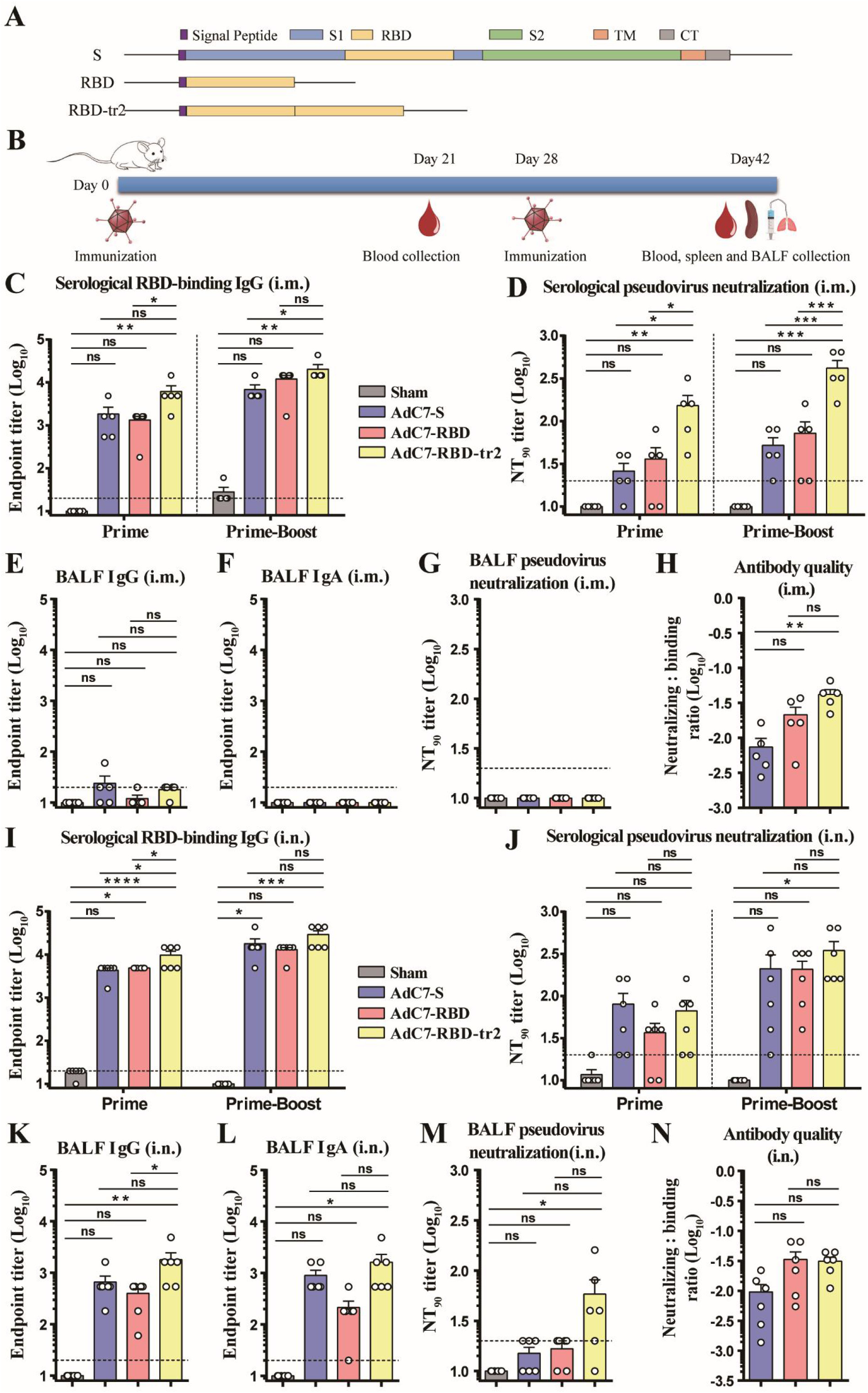
Characterization of the humoral immune responses of BALB/c mice immunized with AdC7 vaccines. (A) Schematic demonstration of antigen constructs of full-length S, RBD and RBD-tr2. Sequences encoding signal peptide was original from MERS-CoV S protein. (B) Schedule of animal experiments. Female BALB/c mice (6-8-weeks old) were immunized with two doses of 1 × 10^11^ vp of AdC7 vaccines through the i.m. or i.n. route, respectively. AdC7-empty was inoculated as a sham vaccine. Sera were collected three weeks post prime immunization. BALF, sera and spleen were collected two weeks post boost immunization. (C - H) Antibody responses of BALB/c mice (n = 5) vaccinated with AdC7 vaccines via the i.m. route. (C) Measurement of SARS-CoV-2 monomeric RBD-binding IgG endpoint titers of serum samples from mice immunized via the i.m. route. Prime indicates serum samples collected at day 21 post the first dose vaccination. Prime-Boost indicates serum samples collected at day 14 post the second dose vaccination. (D) Measurement of SARS-CoV-2 pseudovirus NT_90_ of serum samples from mice immunized via the i.m. route. (E - G) Measurement of SARS-CoV-2 monomeric RBD-binding IgG (E) and IgA (F) endpoint titers and pseudovirus NT_90_ (G) of BALF from mice immunized via the i.m. route. (H) Antibody quality of serum samples from two-dose immunized mice via the i.m. route. The ratio is the pseudovirus neutralization titer : S protein-binding IgG titer. NT_90_ and S protein-binding titers were shown in Fig. 1D and Fig. S3A. (I - N) Antibody responses of BALB/c mice (n = 6) vaccinated with AdC7 vaccines via the i.n. route. (I) Measurement of SARS-CoV-2 monomeric RBD-binding IgG endpoint titers of serum samples from mice immunized via the i.n. route. (J) Measurement of SARS-CoV-2 pseudovirus NT_90_ of serum samples from mice immunized via the i.n. route. (K - M) Measurement of SARS-CoV-2 monomeric RBD-binding IgG (K) and IgA (L) endpoint titers and pseudovirus NT_90_ (M) of BALF from mice immunized via the i.n. route. (N) Antibody quality of serum samples from two-dose immunized mice via the i.n. route. The ratio is the pseudovirus neutralization titer : S protein-binding IgG titer. NT_90_ and S protein-binding titers were shown in Fig. 1J and Fig. S3B. Data are means ± SEM (standard errors of means). *P* values were analyzed with one-way ANOVA (ns, *P* > 0.05; *, *P* < 0.05; **, *P* < 0.01; ***, *P* < 0.001; ****, *P* < 0.0001). The dashed line indicates the limit of detection.

To evaluate the immunogenicity of the recombinant AdC7 vaccines, groups of BALB/c mice (n = 5) were immunized on day 0 and 28 with 1 × 10^11^ vp of AdC7 vaccines via intramuscular (i.m.) injection (Fig. 1B). AdC7 without any transgene (AdC7-empty) was injected as the sham control. Blood samples were collected on day 21 and 42 (Fig. 1B). The serological RBD-binding IgG and IgA titers was measured. RBD-specific IgG were robustly induced by AdC7-S, AdC7-RBD and AdC7-RBD-tr2 (Fig. 1C), while RBD-specific IgA antibodies were only moderately induced in the mice vaccinated with AdC7-RBD-tr2 (fig. S2A). As neutralizing antibodies play a crucial role in protection, pseudovirus displaying the SARS-CoV-2 S protein was used to detect the neutralizating antibody titers in the sera of immunized mice. We found 90% neutralization titer (NT_90_) of the mice sera were increased post boost vaccination for each AdC7 vaccine (Fig.1D). Two doses of AdC7-S, AdC7-RBD and AdC7-RBD-tr2 elicited average NT_90_ titers of 52, 72 and 416, respectively (Fig. 1D). AdC7-RBD-tr2 showed significantly higher neutralizing titers compared to the other constructs (Fig. 1D).

SARS-CoV-2 was transmitted through respiratory droplets and caused lung damage diseases. Specific immune response in the air-way and lung would confer protection against SARS-CoV-2 infection in situ. Immunized mice were euthanized for bronchoalveolar lavage fluid (BALF) collection. RBD-binding IgG, IgA and pseudovirus NT_90_ were measured for the BALF. As expected, a small amount of RBD-binding IgG was detected in these AdC7 vaccines without significant difference compared with sham vaccine (Fig. 1E). Neither RBD-binding IgA nor neutralizing antibodies can be detected in the BALF of mice immunized with AdC7-S, AdC7-RBD or AdC7-RBD-tr2 (Fig. 1, F and G). In order to compare the quality of antibody, the serological neutralizing : binding ratio were calculated for these three constructs. The results showed AdC7-RBD-tr2 elicited highest neutralizing : binding ratio, suggesting its advantage of immunofocusing that potentially reduced ADE risk (Fig. 1H and S3A).

As low levels of RBD-binding and neutralizing antibodies were detected in the BALF of mice immunized with each of the AdC7 vaccines via the i.m. route, we sought to immunize BALB/c mice (n = 6) intranasally (i.n.) to strengthen mucosal immunity. On day 0 and 28, the BALB/c mice were vaccinated with 1 × 10^11^ vp of AdC7 via the i.n. route. On day 21 and 42, serum samples were collected for antibody titration (Fig. 1B). AdC7-S, AdC7-RBD and AdC7-RBD-tr2 all elicited substantial RBD-binding IgG and IgA (Fig. 1I and fig. S2B). Two-dose immunizations of AdC7-S, AdC7-RBD and AdC7-RBD-tr2 induced the pseudovirus NT_90_ titers of 210, 207 and 347, respectively (Fig 1J). The BALF of mice were also collected at day 42 for further evaluation. High levels of both RBD-binding IgG and IgA were detected in the BALF of the mice immunized with AdC7-S, AdC7-RBD or AdC7-RBD-tr2 (Fig. 1, K and L). The average endpoint titers of RBD-binding IgG and IgA of AdC7-RBD-tr2 group were 1800 and 1620, respectively, with both values higher than the other two constructs (Fig. 1, K and L). Moreover, pseudovirus NT_90_ of the BALF were assessed, with an average of 10, 15, 17 and 58 for sham, AdC7-S, AdC7-RBD and AdC7-RBD-tr2, respectively (Fig 1M). AdC7-RBD-tr2 group showed higher mucosal pseudovirus NT_90_ compared to either AdC7-S or AdC7 RBD. In addition, we analyzed the antibody quality in mice sera, and demonstrated RBD and RBD-tr2-based vaccines exhibited higher neutralizing : binding ratio than fell-length S-based vaccine (Fig 1N and S3B). In summary, both systemic and mucosal immunity were induced by recombinant vaccines through i.n. vaccination. Besides, AdC7-RBD-tr2 showed the advantage of immunofocusing to induce neutralizing antibodies blocking receptor-binding.

Aside from humoral immunity, cellular immunity also play an important role in protection against the SARS-CoV-2 (*72-76*). To characterize the cellular immune responses induced by the AdC7 vaccines, the same cohort of immunized BALB/c mice were euthanized on day 42 (Fig. 1B). Splenic lymphocytes were harvested and stimulated with an overlapping 11-mer peptide pool spanning the SARS-CoV-2 S protein and analyzed by enzyme-linked immunospot (ELISpot) and intracellular cytokine staining (ICS) assays. IFN-γ ELISpot analysis showed the induction of robust T cell responses for mice vaccinated with AdC7-S, AdC7-RBD or AdC7-RBD-tr2 via both i.m. and i.n. routes (Fig. 2, A and B). Additionally, the flow cytometric analyses showed that all these recombinant AdC7 vaccines induced robust induction of cytotoxic T lymphocytes (CTL), detected as IFNγ and IL-2 production, via both i.m. and i.n. routes. AdC7-RBD-tr2 performed the best (Fig. 2C and 2E). Besides, the CD4+ T cell responses cytokines were moderately induced for all these three constructs (Fig.2D and 2F). In contrast, no substantial Th2 cytokines (IL-4 and IL10) production was detected for all these three constructs (Fig.2D and 2F). These results demonstrated Th1-biased immune responses were induced in mice received AdC7-S, AdC7-RBD or AdC7-RBD-tr2 vaccines by both i.m. and i.n. routes.

**Fig. 2.**
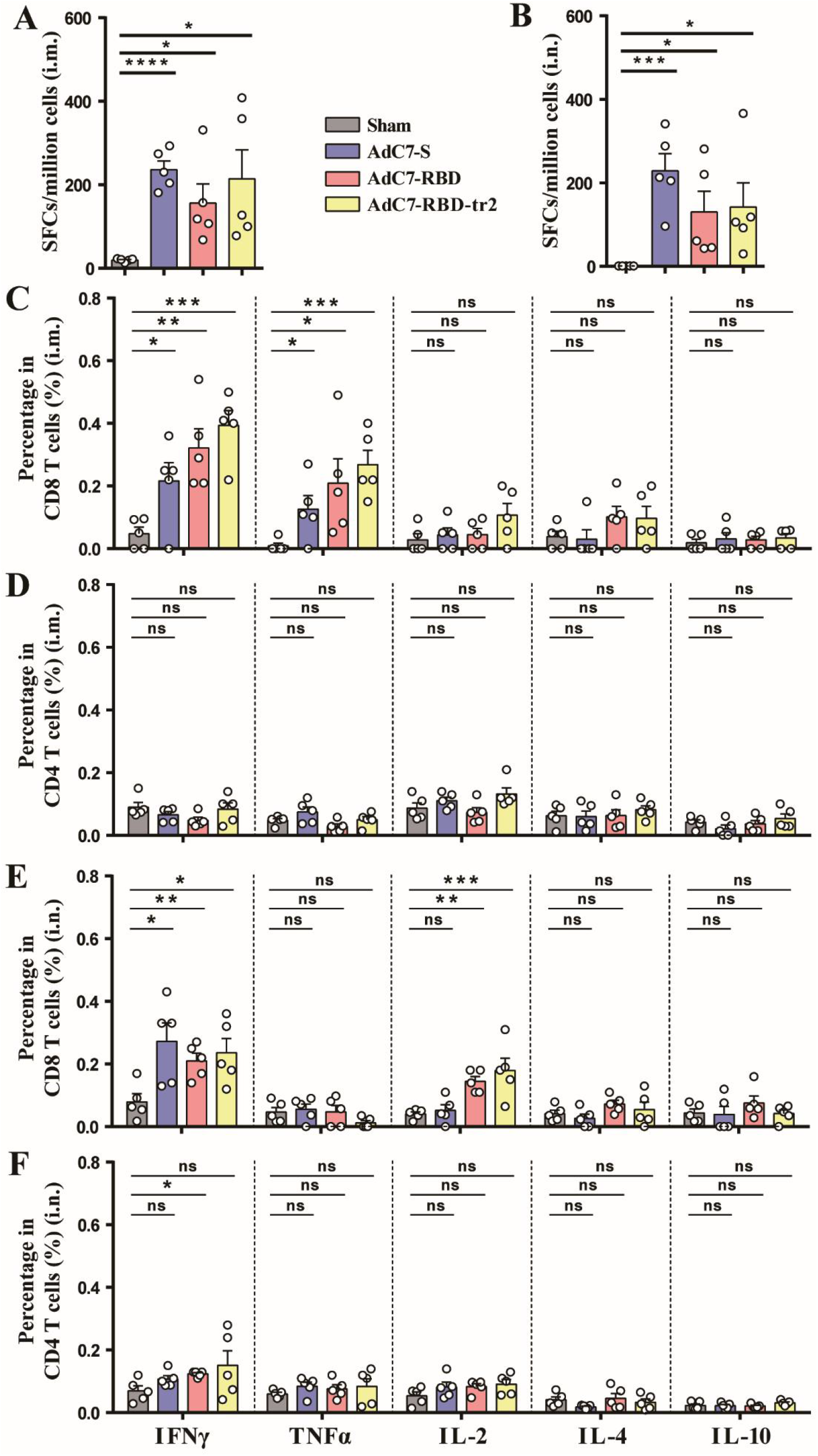
Characterization of the cellular immune responses. BALB/c mice were immunized with 1 × 10^11^ vp AdC7-S, AdC7-RBD, AdC7-RBD-tr2 or Sham (AdC7-empty) as described in Fig. 1B. Mice splenocytes were isolated and analyzed by ELISpot and ICS assays. (A and B) ELISpot assays were performed to evaluation of the IFNγ secretion of splenocytes after SARS-CoV-2 S peptides stimulation for mice immunized through the i.m. (A) and i.n. (B) route. (C - F) ICS assays were conducted to analyze the CD8+ and CD4+ T cell responses of mice immunized via the i.m. (C and D) and i.n. (E and F) route. (C and D) Quantification of the frequency of IFNγ-, TNFα-, IL-2-, I-4- and IL-10-producing CD8+ T cells (C) and CD4+ T cells (D) of splenocytes from mice with i.m. immunization. (E and F) Quantification of the frequency of IFNγ-, TNFα-, IL-2-, I-4- and IL-10-producing CD8+ T cells (E) and CD4+ T cells (F) of splenocytes from mice with i.n. immunization. Data are means ± SEM. *P* values were analyzed with *t* test (ns, *P* > 0.05; *, *P* < 0.05; **, *P* < 0.01; ***, *P* < 0.001; ****, *P* < 0.0001).

Given the fact that AdC7-RBD-tr2 elicited the most robust RBD-binding IgG, IgA, neutralizing antibodies and CTLs, AdC7-RBD-tr2 vaccine was chosen for the further protection efficacy against SARS-CoV-2 challenge. Since wild type BALB/c mice are not sensitive to SARS-CoV-2 infection due to the low binding affinity between mouse ACE2 and S protein, we transduced the recombinant Ad5 expressing human ACE2 (Ad5-hACE2) into BALB/c mouse lung via i.n. route to rapidly generate a mouse model (*77*). Five days later, the vaccinated mice were challenged with 5 × 10^5^ TCID_50_ SARS-CoV-2 (Fig. 3A). AdC7-RBD-tr2 inoculation via i.n. route induced BALB/c mice producing substantial titers of RBD-binding IgG, IgA (Fig. 3, B and C) and pseudovirus neutralizing antibody (Fig. 3D). Additionally, live virus neutralizing assay revealed a mean titer of 120 (one-dose) and 154 (two-dose), respectively, upon AdC7-RBD-tr2 vaccination (Fig. 3E). Three days post challenge, lungs of mice were harvested for virus titration and pathologic analysis. High levels of viral RNA were observed with a mean titer of 9.83 log_10_ RNA copies/g in mice lung in the one dose sham group (Fig. 3F). Whereas low level of viral RNA was observed in the AdC7-RBD-tr2-vaccinated mice with a mean titer of 6.00 log_10_ RNA copies/g (Fig. 3F). In the prime-boost groups of mice, the mean titer of viral RNA (log_10_) in mice lung per gram was 9.47 and 5.46 for Sham and AdC7-RBD-tr2, respectively (Fig. 3F). Because a fraction of viral RNA in the lung were probably originated from input challenge virus, levels of subgenomic RNA (sgRNA) were also measured to quantify the live virus. The sgRNA were generated in the infected cells during virus replication but were absent in the virions (*78*). As a result, high levels of sgRNA were observed with a mean titer of 8.87 and 8.90 log_10_ RNA copies/g in sham group with the one-dose and two-dose regimens, respectively. Encouragingly, viral sgRNA could not be detected in any lung samples from both one or two doses of AdC7-RBD-tr2 vaccinated mice (Fig 3G), indicating the complete protection efficacy of AdC7-RBD-tr2 vaccine.

**Fig. 3.**
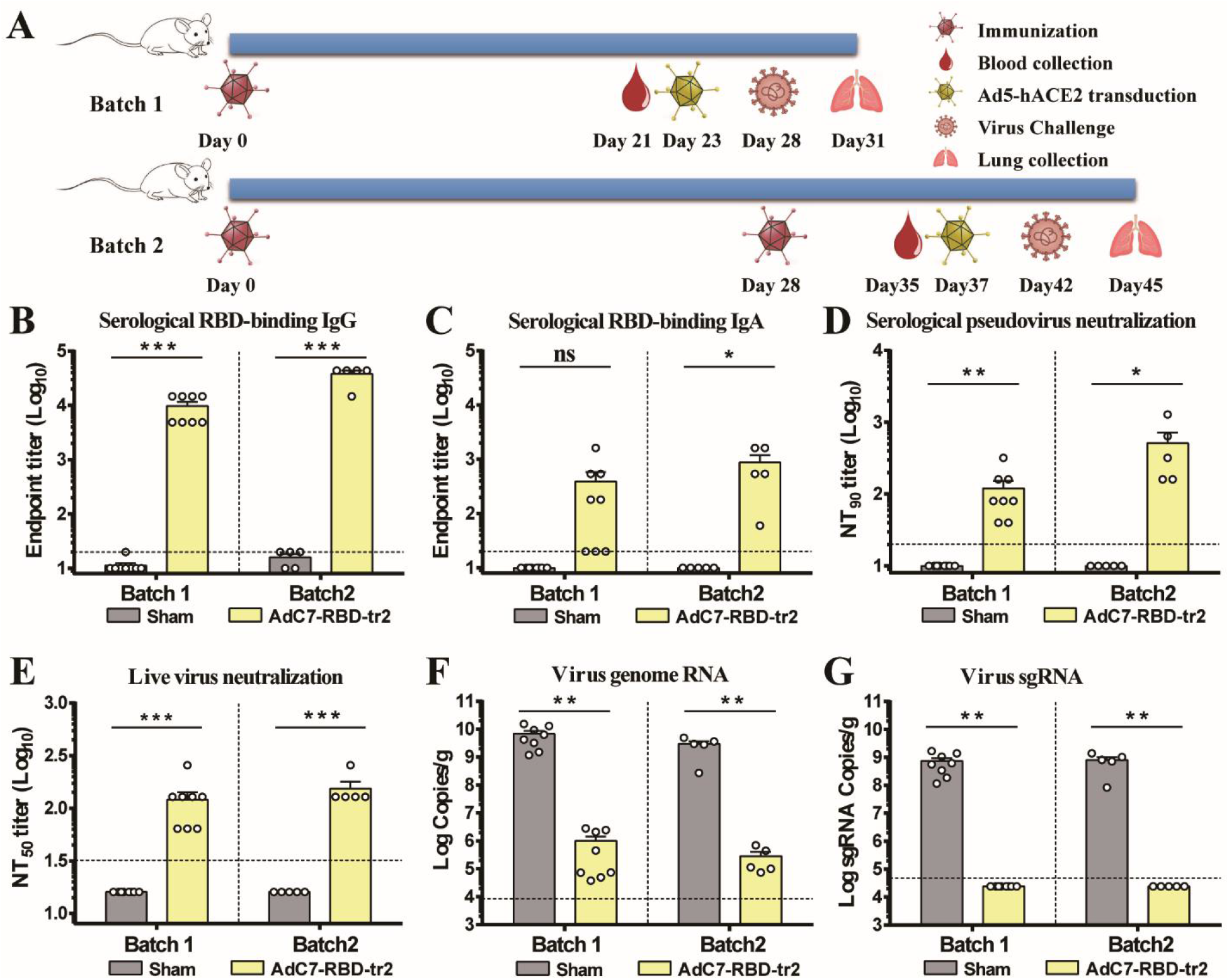
Protective efficacy of AdC7-RBD-tr2 against SARS-CoV-2. (A) Immunization and challenge schedule. Female BALB/c mice (6-8-weeks old) of batch 1 (n = 8) and batch 2 (n = 5) received one-dose or two-dose of 1 × 10^11^ vp of AdC7-RBD-tr2 through the i.n. route, respectively. The same dose of sham vaccine (AdC7-empty) was i.n. infected as the control. Prior to SARS-CoV-2 challenge, blood samples from vaccinated mice were collected for antibodies titration. At five days post Ad5-hACE2 transduction, mice were i.n. challenged with 5 × 10^5^ TCID_50_ SARS-CoV-2. Animals were euthanized and necropsied on 3 dpi. and lung tissues were harvested for virus titration and pathological examination. (B and C) Measurement of SARS-CoV-2 monomeric RBD-binding IgG (B) and IgA (C) endpoint titers of sera. (D) Measurement of pseudovirus NT_90_ of sera. (E) Measurement of real SARS-CoV-2 neutralizing antibody titers (NT_50_) of sera. (F and G) SARS-CoV-2 titration from lung tissues by qRT-PCR probing virus genome RNA (F) and sgRNA (G). Data are means ± SEM. *P* values were analyzed with *t* test (ns, *P* > 0.05; *, *P* < 0.05; **, *P* < 0.01; ***, *P* < 0.001). The dashed line indicates the limit of detection.

In order to further evaluate vaccine protection against lung damage by SARS-CoV-2, histopathological analysis was performed. The results exhibited mice from sham group developed apparent viral pneumonia characterized by thickened alveolar walls, vascular congestion and inflammatory cell infiltration (Fig. 4, A, B, I and J), whereas a marked attenuation of pathological damage and inflammatory response were seen in the lung tissues of mice vaccinated with AdC7-RBD-tr2 (Fig. 4, C, D, K and L). More importantly, immunofluorescence analysis of lung section stained with anti-SARS-CoV-2 NP antibody revealed the virus presence in the lung of sham group (Fig. 4, E, F, M and N), but not in that of AdC7-RBD-tr2 group (Fig. 4, G, H, O and P). In summary, these results demonstrated the AdC7-RBD-tr2 vaccine robustly elicited mice immune responses against SARS-CoV-2 and confer nearly sterilizing immunity against SARS-CoV-2 infection.

**Fig. 4.**
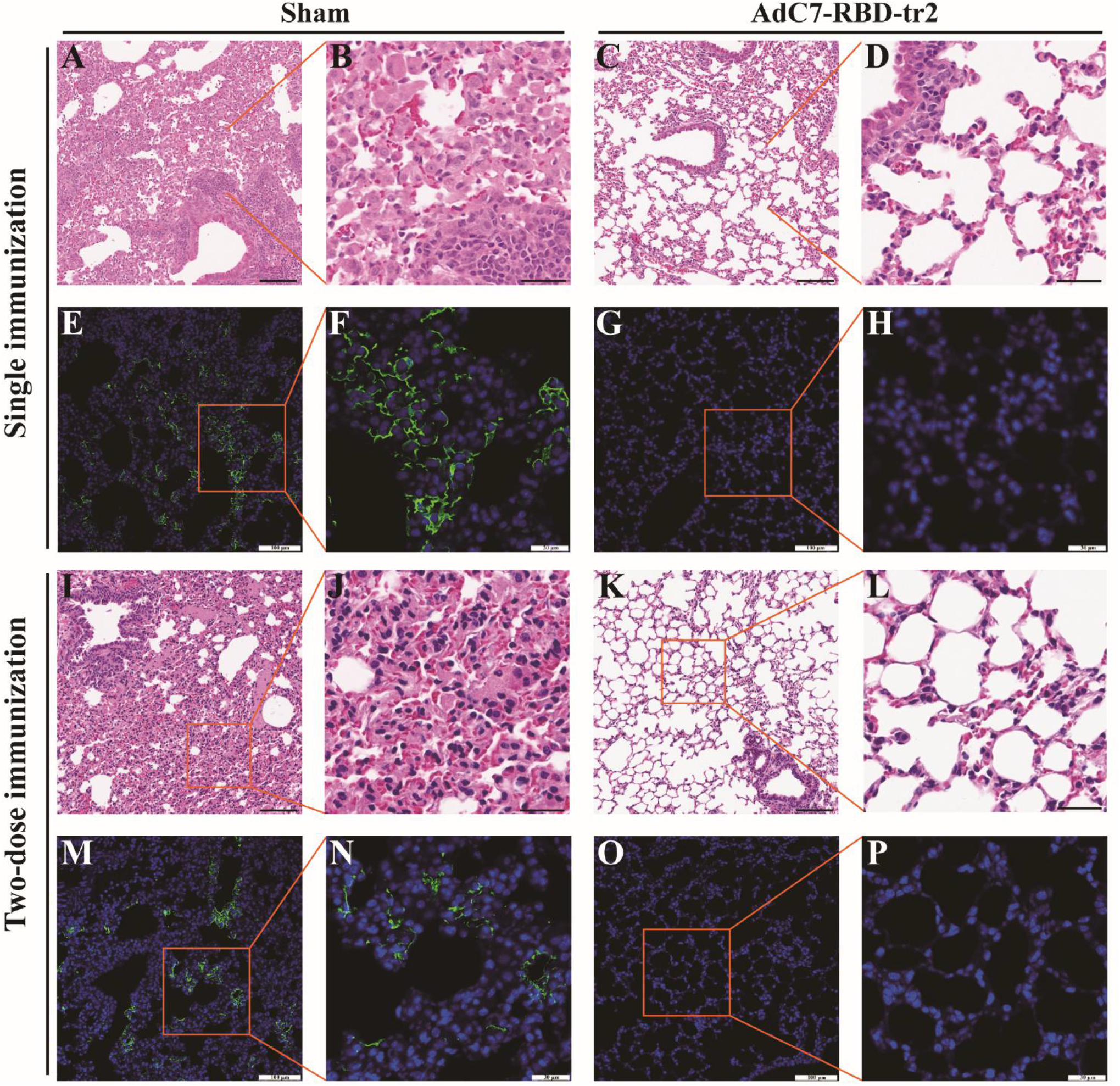
Protection against lung infection and lesion by AdC7-RBD-tr2. Mice lung tissues were fixed in 4% paraformaldehyde, embedded in paraffin, and then sectioned. Tissue sections (4 μm) were stained with H&E or anti-SARS-CoV-2 nucleoprotein antibody for pathological examination and virus probing. (A - H) Histopathology and immunofluorescence analysis of lung tissue sections from batch 1 mice with single immunization. (I - P) Histopathology and immunofluorescence analysis of lung tissue sections from batch 2 mice with two doses immunization. (A - D, and I - L) Images of lung pathology from sham group (A, B, I and J) and AdC7-RBD-tr2 group (C, D, K, and L). Both low magnifications (A, C, I and K) and high magnifications (B, D, J and L) are shown. (E - H, and M - P) Images of immunofluorescence from sham group (E, F, M and N) and AdC7-RBD-tr2 group (G, H, O, and P). Both low magnifications (E, G, M and O) and high magnifications (F, H, N and P) are shown. Scale bar in low magnifications images, 100 μm. Scale bar in high magnifications images, 30 μm.

## Discussion

Major safety concerns for the vaccine include the vaccine-associated enhanced respiratory disease (ERD), which is associated with unbalanced Th2-biased T cell responses, characterized by enhanced inflammation and immunopathology, and ADE, which is caused by virus uptake into lymphocyte expressing Fc receptor, resulting in increased viral replication and antibody Fc-mediated effector functions (*79-81*). ADE has been reported for SARS, MERS and other human respiratory virus infections including RSV and measles, which raises a potential concern for the risk of vaccine-enhanced COVID-19 severity (*82-86*). Although no convincing evidence demonstrated a role for ADE in human COVID-19 pathology, the risk from vaccine immunization should be closely pay attention during vaccine designation (*59*).

Non-neutralizing antibodies or antibodies at sub-neutralizing concentrations bind to viral antigens without blocking or clearing infection was thought to be a major reason for ADE. Up to now, all reported adenovirus vector vaccines against COVID-19 were designed to express full length S protein or modified S protein(*19, 20, 24, 32, 87*). Specific antibody could be elicited targeting the non-neutralization epitopes on S protein with binding activity. In order to decrease the potential risk of ADE, we constructed a recombinant AdC7 vaccine expressing RBD-tr2 protein and demonstrated it induced robust humoral responses with high antibody quality and Th1-biased T cell responses. In addition, mucosal immunity could also be effectively activated through i.n. immunization, characterized by amounts of RBD-binding IgA and neutralization antibody detected in BALF. Compared with the full-length S construct, RBD-tr2 antigen may induce less non-neutralizing antibodies and enhance antibody quality. The overall immunity elicited by AdC7-RBD-tr2 vaccine confer mice complete protection against COVID-19 without infection enhancement or immunopathological exacerbation. Collectively, these results suggest AdC7-RBD-tr2 is a promising candidate vaccine to move forward for preclinical and clinical trials for prophylaxis of COVID-19.

## Acknowledgement

We are grateful to T. Zhao [Institute of Microbiology, Chinese Academy of Sciences (CAS)] for her technical assistance with flow cytometry experiments. We thank X. Zhang (Institute of Microbiology, CAS) for her technical assistance with using laser scanning confocal microscope. We thank G. Wong (Institut Pasteur of Shanghai, CAS) for providing us the Ad5-hACE2 virus. We thank the staff of ABSL-3 at Institute of Microbiology, CAS for assistance about the work operated in ABSL-3.

## Author contributions

K.X., G.F.G., and L.D. initiated and coordinated the project. Q.W., W.H., Q.X., G.F.G., and L.D. supervised the project. K.X. and L.D. designed the experiments. K.X., Y.A., Y.H., T.Z., C.L. P.G., and S.X. conducted the experiments. Q.L., F.F., and H.L. discussed the experiments. W.H. and Y.W. provided the SARS-CoV-2 pseudovirus. W.J.L. designed the SARS-CoV-2 peptides and provided advices for cellular immunity evaluation. Y.B. supervised the experiments with SARS-CoV-2 in ABSL-3. K.X., Q.X., G.F.G., and L.D. analyzed the data. K.X., G.F.G., and L.D. wrote the manuscript. Q.L., W.J.L., Y.W., D.Z., Q.W., W.H., Q.X., G.F.G., and L.D. review and discussed the manuscript. K.X., G.F.G., and L.D. edited the manuscript.

## Materials and Methods

### Materials

#### Cells, viruses, and animals

Human embryonic kidney 293 (HEK293) cells (ATCC CRL-1573), HEK293T cells (ATCC CRL-3216), Huh7 hepatoma cells (Institute of Basic Medical Sciences, CAMS) and VERO-E6 were all maintained in complete Dulbecco’s modified Eagle’s medium (DMEM, Invitrogen, USA) supplemented with 10% fetal bovine serum (FBS) and incubated at 37°C under 5% CO_2_.

SARS-CoV-2 was isolated from an infected patient in Institute of Microbiology, Chinese Academy of Science (IMCAS), Beijing, China, which were further propagated in VERO-E6 cells and titrated by tissue culture infectious dose 50 (TCID_50_) assay on VERO-E6 cells. Pseudovirus displaying SARS-CoV-2 S protein was kindly provided by Youchun Wang (Institute for Biological Product Control, National Institutes for Food and Drug Control, Beijing, China).

Specific pathogen-free (SPF) 6-8-weeks old female BALB/c mice were purchased from Beijing Vital River Laboratory Animal Technology Co., Ltd. (licensed by Charles River), and housed under SPF conditions in the laboratory animal facilities at IMCAS. All animals were allowed free access to water and standard chow diet and provided with a 12-hour light and dark cycle.

### Ethics statement

All animal experiments were approved by the Committee on the Ethics of Animal Experiments of the IMCAS, and conducted in compliance with the recommendations in the Guide for the Care and Use of Laboratory Animals of the IMCAS Ethics Committee.

## Methods

### Construction and production of recombinant chimpanzee adenovirus

An E1- and E3-deleted, replication-deficient recombinant chimpanzee type 7 adenovirus (AdC7) vector was used to construct recombinant AdC7 vaccines encoding full-length S, RBD or RBD-tr2 of SARS-CoV-2 (GenBank accession number YP_009724390). The full-length S construct contains MERS S protein signal peptide (MIHSVFLLMFLLTPTES) and amino acids 16 to 1273 of the S protein of SARS-CoV-2. The RBD construct contains the same signal peptide and amino acids 319 to 537 of the S protein of SARS-CoV-2. The RBD-tr2 construct contains the same signal peptide and two RBD (amino acids 319 to 537 of the S protein) connected as tandem repeat without any linker sequence. The cassette of full-length S, RBD and RBD-tr2 were cloned into pAdC7, forming recombinant adenovirus genome, respectively. These recombinant adenovirus genomes were linearized and transfected into HEK293 cells to rescue the recombinant adenovirus, which was further propagated and purified by cesium chloride density gradient centrifugation as previously described (*88*).

### Western blot

HEK 293T cells were preplated in 6-well plate, followed by infected with 1 × 10^9^ vp of AdC7-S, AdC7-RBD, AdC7-RBD-tr2 or AdC7-empty as sham control. Forty-eight hours post infection, cells were lysed, and culture supernatants were collected. Protein samples were separated by 12% SDS-PAGE and analyzed by Western blotting with rabbit anti-RBD of SARS-CoV-2 polyclonal antibody. Goat anti-rabbit IgG-horseradish peroxidase (HRP) antibodies were used as secondary antibodies. The membranes were developed by SuperSignal West Pico chemiluminescent substrate (Thermo Fisher Scientific, USA).

### Proteins expression and purification

Monomer RBD protein of SARS-CoV-2 was expressed and purified as previously described (*13*). Briefly, signal peptide sequence of MERS-CoV S protein (MIHSVFLLMFLLTPTES) was added to the RBD protein (S protein 319-541, GenBank: YP_009724390) N terminus for protein secretion, and a hexa-His tag was added to the C terminus to facilitate further purification processes. The coding sequence was codon-optimized for mammalian cell expression and synthesized by GENEWIZ, China. Then, the construct was cloned into the pCAGGS vector and transiently transfected into HEK 293T cells. After 3 days, the supernatant was collected and soluble protein was purified by Ni affinity chromatography using a HisTrap™ HP 5 mL column (GE Healthcare). The sample was further purified via gel filtration chromatography with HiLoad® 16/600 Superdex® 200 pg (GE Healthcare) in a buffer composed of 20 mM Tris-HCl (pH 8.0) and 150 mM NaCl.

### Immunization

AdC7-S, AdC7-RBD and AdC7-RBD-tr2 was diluted in PBS. Empty vector AdC7 (AdC7-empty) was used as a sham control. Female BALB/c mice at 6 to 8 weeks of age were immunized with 1 × 10^11^ vp of vaccine or sham control through the i.m. or i.n. route. The second dose was as same as the first dose and given 28 days post prime vaccination. The sera were collected as indicated in figures legends.

### Enzyme-linked immunosorbent assay (ELISA)

Binding properties of murine sera to monomer RBD or S protein were determined by ELISA. 96-well plates (3590; Corning, USA) were coated over-night with 3 μg/ml of monomer RBD or S protein (Sino Biological, China) in 0.05 M carbonate-bicarbonate buffer, pH 9.6, and blocked in 5% skim milk in PBS. Serum or BALF samples were serially diluted and added to each well. Plates were incubated with goat anti-mouse IgG-HRP antibody or goat anti-mouse IgA-HRP antibody and subsequently developed with 3,3’,5,5’-tetramethylbenzidine (TMB) substrate. Reactions were stopped with 2 M hydrochloric acid, and the absorbance was measured at 450 nm using a microplate reader (PerkinElmer, USA). The endpoint titers were defined as the highest reciprocal dilution of serum to give an absorbance greater than 2.5-fold of the background values. Antibody titer below the limit of detection was determined as half the limit of detection.

### Pseudovirus neutralization assay

SARS-CoV-2 pseudovirus preparation and neutralization assay were carried out by a previously published method (*89*), with some modifications. Briefly, mice serum or BALF samples were 2-fold serially diluted and incubated with an equal volume of 100 TCID_50_ pseudovirus at 37°C for 1 hour. The medium was also mixed with pseudovirus as control. Then the mixture was transferred to pre-plated Huh7 cell monolayers in 96-well plates. After incubation for 24 hours, the cells were lysed and luciferase activity was measured by the Luciferase Assay System (Promega, USA) according to the manufacturer’s protocol. NT_90_ was defined as the highest reciprocal serum dilution at which the relative light units (RLUs) were reduced by greater than 90% compared with virus control wells. NT_90_ below the limit of detection was determined as half the limit of detection.

### Live SARS-CoV-2 neutralization assay

The neutralizing activity of mouse serum was assessed using a previously described SARS-CoV-2 neutralization assay (*13*). Briefly, sera from immunized mice were 4-fold serially diluted and mixed with the same volume of SARS-CoV-2 (100 TCID_50_), incubated at 37°C for 1 hour. Thereafter, 100 μL virus-serum mixture was transferred to pre-plated VERO-E6 cells in 96-well plates. Inoculated plates were incubated at 37°C for an additional 72 hours, following which the cytopathic effect was observed microscopically. The neutralization titers were defined as the reciprocal of serum dilution required for 50% neutralization of viral infection. All the live virus neutralization assay was conducted under biosafety level 3 (BSL3) facility in IMCAS.

### ELISpot assay

To detect antigen-specific T lymphocyte responses, an IFNγ-based ELISpot assay was performed as previously described (*88*), with some modifications. Briefly, an S peptide pool consisting of 15-18-mers (overlapping by 11 amino acids) and spanning the entire S protein of SARS-CoV-2 were synthesized. Spleens of vaccinated BALB/c mice were harvested at 2 weeks post the second dose immunization and plenocytes were isolated. Flat-bottom, 96-well plates were precoated with 10 μg/ml anti-mouse IFNγ Ab (BD Biosciences, USA) overnight at 4°C and then blocked for 2 hours at 37°C. Mouse splenocytes were added to the plate. Then, the peptide pool (2 μg /ml individual peptide) was added to the wells. Phytohemagglutinin (PHA) was added as a positive control. Cells incubated without stimulation were employed as a negative control. After 24 h of incubation, the cells were removed, and the plates were processed in turn with biotinylated IFNγ detection antibody, streptavidin-HRP conjugate, and substrate. When the colored spots were intense enough to be visually observed, the development was stopped by thoroughly rinsing samples with deionized water. The numbers of the spots were determined using an automatic ELISpot reader and image analysis software (Cellular Technology Ltd.).

### ICS and flow cytometry

ICS assays were performed as previously described (*90*), with some modifications. Briefly, mouse splenocytes were added to the plate (2 × 10^6^/well) and then stimulated with the peptide pool (2 μg /ml for individual peptide) for 5 h. The cells were incubated with GolgiStop (BD Biosciences, USA) for an additional 6 h at 37°C. Then, the cells were harvested and stained with anti-CD3 (BioLegend), anti-CD4 (BioLegend) and anti-CD8α (BioLegend) surface markers. The cells were subsequently fixed and permeabilized in permeabilizing buffer (BD Biosciences, USA) and stained with anti-mouse anti-IFNγ (BioLegend), anti-TNFα (BioLegend), anti-IL-2 (BioLegend), anti-IL-4 (BioLegend) and anti-IL-10 (BioLegend) antibodies. All fluorescent lymphocytes were gated on a FACSAria flow cytometer (BD Biosciences, USA).

### Animal protection against virus challenge

To evaluate the protection efficacy of vaccine candidates against SARS-CoV-2, a recombinant adenovirus Ad5-hACE2 transducing BALB/c mice model was used. Immunized BALB/c mice were i.n infected with 8 × 10^8^ vp of Ad5-hACE2. Five days later, the transduced mice were challenged with 5 × 10^5^ TCID_50_ of SARS-CoV-2 via the i.n. route. Three days post challenge, mice were euthanized and necropsied. Lung tissues were collected and split into two parts for virus titration and pathological examination. All animal experiments with SARS-CoV-2 challenge were conducted under animal biosafety level 3 (ABSL3) facility in IMCAS.

### qRT-PCR

Mice lung tissues were weighed and homogenized. Virus RNA was isolated from 50-μl supernatants of homogenized tissues using a nucleic acid extraction instrument MagMAX™ Express Magnetic Particle Processor (Applied Biosystems, USA). SARS-CoV-2-specific quantitative reverse transcription-PCR (qRT-PCR) assays were performed using a FastKing One Step Probe RT-qPCR kit (Tiangen Biotech, China) on a CFX96 Touch real-time PCR detection system (Bio-Rad, USA) according to the manufacturer’s protocol. Two sets of primers and probes were used to detect a region of the N gene of viral genome (*90*) and a region of E gene of sgRNA of SARS-CoV-2 (*91*), respectively, with sequences as follows:

N-F, GACCCCAAAATCAGCGAAAT;

N-R, TCTGGTTACTGCCAGTTGAATCTG;

N-probe, FAM-ACCCCGCATTACGTTTGGTGGACC-TAMRA (where FAM is 6-carboxyfluorescein, and TAMRA is 6-carboxytetramethylrhodamine);

sgRNA-E-F, CGATCTCTTGTAGATCTGTTCTC;

sgRNA-E-R, ATATTGCAGCAGTACGCACACA;

sgRNA-E-probe, FAM-ACACTAGCCATCCTTACTGCGCTTCG-TAMRA.

Viral loads were expressed on a log10 scale as viral copies/gram after calculation with a standard curve. Viral copy numbers below the limit of detection were set as half the limit of detection.

### Histopathology analysis

Mice lung tissues were fixed in 4% paraformaldehyde, dehydrated, embedded in paraffin, and then sectioned. Tissue sections (4 μm) were deparaffinized in xylene and stained with hematoxylin and eosin (H&E) for pathological examination, such as peribronchiolitis, interstitial pneumonitis and alveolitis. Besides, tissue sections were stained with anti-SARS-CoV-2 nucleoprotein antibody (Sino Biological, China) to detect virus infection.

### Data analysis

Data are expressed as the means ± standard errors of the means (SEM). For all analyses, *P* values were analyzed by one-way ANOVA with multiple comparisons or unpaired *t* test. Graphs were generated with GraphPad Prism software.

## Supplementary Materials for

**Fig. S1.**
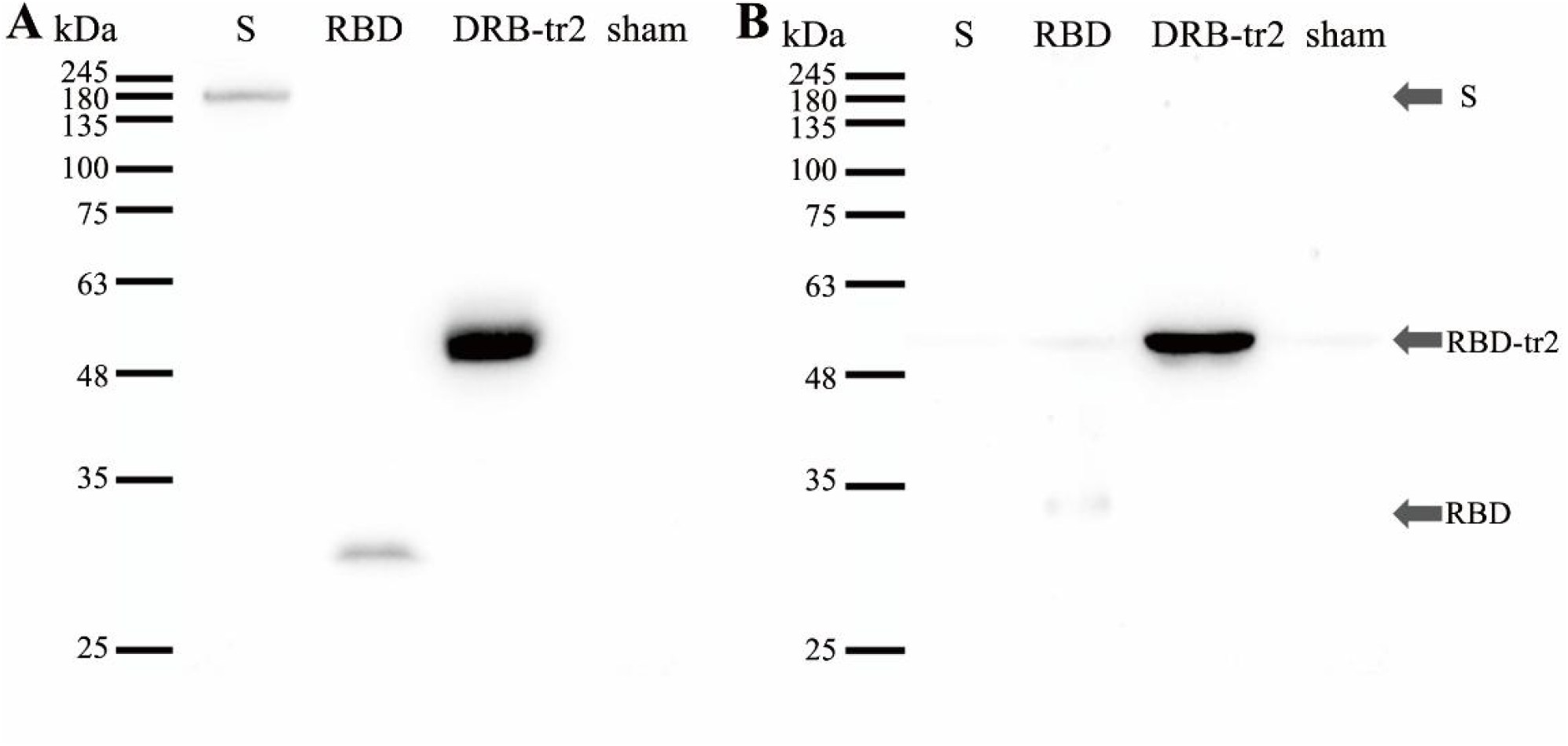
Analysis of transgene expression by western blot. HEK293T cells were infected with AdC7-S, AdC7-RBD, AdC7-RBD-tr2 or AdC7-empty (sham). Antigen proteins were probed in cell lysates (A) and supernatants (B).

**Fig. S2.**
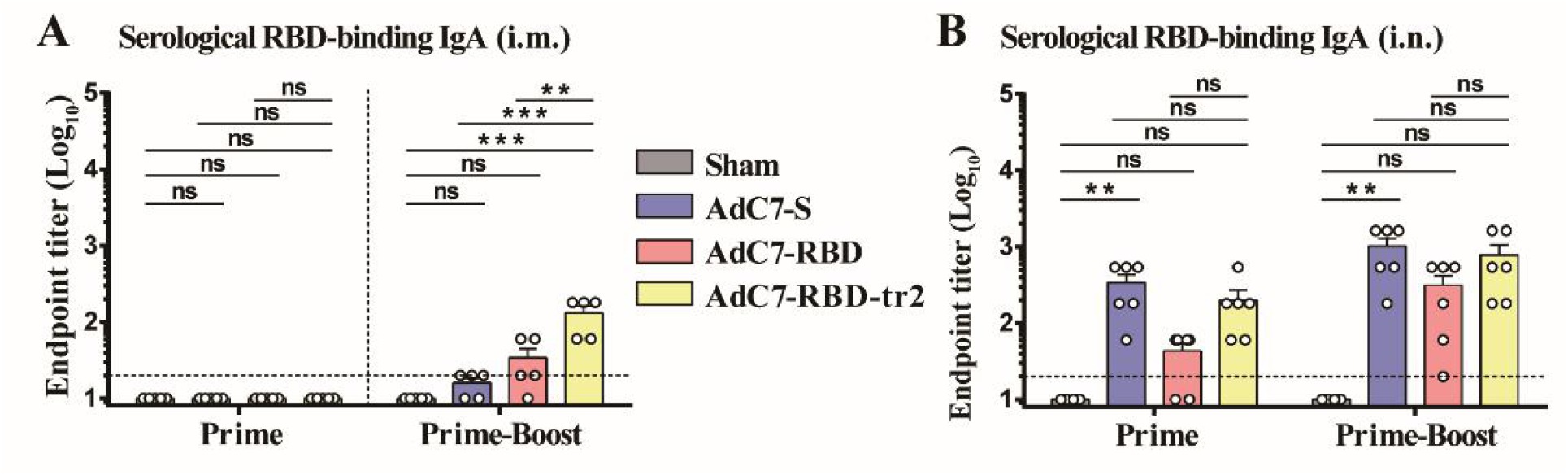
Induction of RBD-binding IgA in BALB/c mice immunized with AdC7 vaccines. Measurement of SARS-CoV-2 monomeric RBD-binding IgA endpoint titers of serum samples from mice immunized via the i.m. (A) and i.n. (B) route. Prime indicates serum samples collected at day 21 post the first dose vaccination. Prime-Boost indicates serum samples collected at day 14 post the second dose vaccination. Data are means ± SEM. *P* values were analyzed with one-way ANOVA (ns, *P* > 0.05; **, *P* < 0.01; ***, *P* < 0.001). The dashed line indicates the limit of detection.

**Fig. S3.**
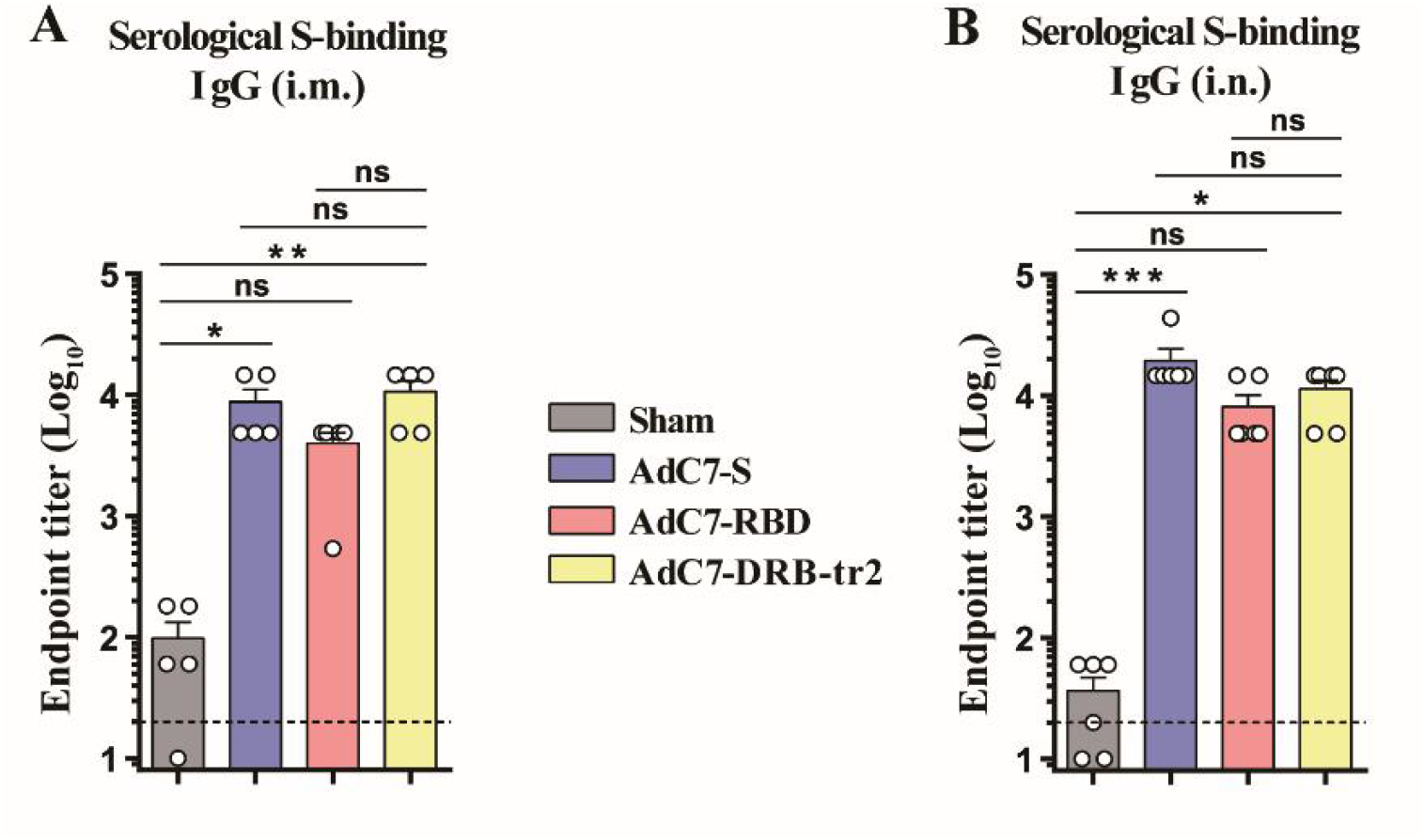
Induction of SARS-CoV-2 S protein-binding IgG in BALB/c mice immunized with AdC7 vaccines. Measurement of SARS-CoV-2 full-length S protein-binding IgG endpoint titers of serum samples from mice immunized two doses of AdC7 vaccines via the i.m. (A) and i.n. (B) route. Data are means ± SEM. *P* values were analyzed with one-way ANOVA (ns, *P* > 0.05; *, *P* < 0.05; **, *P* < 0.01; ***, *P* < 0.001). The dashed line indicates the limit of detection.

